# Beyond model-free Pavlovian responding: a two-stage Pavlovian-instrumental transfer paradigm

**DOI:** 10.64898/2026.03.06.710018

**Authors:** Laura A. Wirth, Nassim Sadedin, Björn Meder, Daniel J. Schad

**Affiliations:** Institute for Mind, Brain and Behavior, HMU Health and Medical University, Potsdam, Germany; Cognitive Sciences, University of Potsdam, Potsdam, Germany

**Author notes:** Corresponding author (LAW).

**Keywords:** Pavlovian conditioning, Pavlovian-instrumental transfer, model-based learning, model-free learning, computational modeling, reinforcement learning, mind wandering

## Abstract

**Background:** Pavlovian responding is a core component of behavior and can be measured via Pavlovian-instrumental transfer (PIT), where Pavlovian responses bias instrumental actions. Standard single-lever PIT paradigms, which assess responses using a single-choice option, cannot dissociate the contribution of model-free versus model-based reinforcement learning. While indirect evidence suggests a role for model-free responding in single-lever PIT, the contribution of model-based strategies is unclear. It also remains unknown whether internal cognitive states, such as mind wandering, impair specifically model-based but not model-free PIT, as is theoretically expected.

**Methods:** We developed a novel, trial-by-trial two-stage PIT paradigm designed to computationally dissociate model-free and model-based Pavlovian responding by leveraging probabilistic state transitions and trial-wise outcome predictions. After each two-stage Pavlovian learning trial, participants performed a single-lever PIT trial as well as a query trial of explicit value judgment. Detailed task instructions were provided to support potential model-based strategies. Computational modeling was used to quantify individual learning strategies. We assessed mind-wandering questionnaires and thought probes.

**Results:** Analysis of query and PIT trials revealed trial-by-trial updating of outcome expectations based on probabilistic task structure, consistent with model-based Pavlovian responding. Behavioral responses during PIT were best explained by a computational model-based reinforcement learning model. In contrast, we found little evidence for model-free Pavlovian responding. Higher levels of mind wandering were associated with reduced model-based control but did not impact model-free indices.

**Conclusion:** We introduce a novel single-lever PIT paradigm that enables fine-grained dissociation of model-free versus model-based Pavlovian response systems. Our findings provide evidence that single-lever PIT can operate through model-based mechanisms, challenging the assumption that single-lever PIT is predominantly model-free. Our findings also indicate that internal attentional states selectively modulate model-based PIT. Given the involvement of Pavlovian responding in numerous psychiatric conditions, our paradigm offers new avenues for understanding maladaptive behavior.

**Author Summary:** Our daily actions are often influenced by cues like the smell of food or the sound of phone notifications that signal potential rewards or losses. These Pavlovian cues can shape our instrumental behavior even though their outcomes do not depend on what we do - a process known as Pavlovian-instrumental transfer (PIT). Here we study the computational learning mechanisms that underlie such PIT effects. While it is often assumed that Pavlovian responding follows simple, automatic rules without a cognitive model of cue consequences (i.e., model-free), evidence also shows a role for cognitive anticipations in Pavlovian responding (i.e., model-based).

In this study, we extend this evidence by showing that PIT responding can be driven by flexible model-based learning. We designed a task to test whether participants use model-free versus model-based strategies to guide PIT, providing detailed task instructions. Using reinforcement learning models, we found that most participants used model-based learning when forming cue-outcome associations. Importantly, people’s attention mattered: when they were more distracted and doing mind wandering, they relied less on model-based strategies.

Our findings suggest that Pavlovian learning is complex, flexible, and influenced by internal mental states, opening new windows to understand decision-making problems in mental health conditions like addiction.

## Introduction

Two computational *reinforcement learning* (RL) mechanisms have been proposed to underlie Pavlovian and instrumental learning and responding [1–3]: *Model-free* learning is based on forming associations between cues, i.e., stimuli or states, and outcomes through trial and error, enabling rapid but inflexible responses that do not require further knowledge of how outcomes arise. In contrast, *model-based* learning involves constructing and utilizing an internal model of the environment that encodes knowledge of state transitions, allowing flexible goal-directed behavior. Distinguishing these mechanisms is critical for understanding how organisms balance efficient habitual actions with flexible goal-directed decision-making [3–6]. Computational models of model-free and model-based learning are well established in instrumental learning [1,7], but remain less explored in Pavlovian learning, which has often been modelled as relying on simple model-free processes [8]. However, recent evidence suggests that Pavlovian learning, and specifically how Pavlovian values bias ongoing instrumental actions, may also engage both model-free and model-based processes, reflecting the dual systems described in RL theory [9].

Key evidence for model-based and model-free Pavlovian responding comes from *Pavlovian Instrumental Transfer* (PIT) tasks. In PIT, Pavlovian cues modulate independently trained instrumental responses, with appetitive cues eliciting approach behavior and aversive cues eliciting avoidance [10–12]. The strength of this influence varies considerably across individuals [13–16] and has been linked to a range of psychiatric symptoms [17]. Importantly, individuals differ not only in the magnitude of PIT effects they exhibit, but also in the type that predominates. These types are classified as general or specific PIT [17,18].

In *general PIT*, Pavlovian cues broadly invigorate or suppress instrumental actions based solely on their motivational valence. For example, a cue predicting one reward (e.g., a monetary win) can enhance responses aimed at obtaining a different reward (e.g., a preferred snack). By contrast, *specific PIT* occurs when Pavlovian cues selectively increase instrumental responses that lead to the same specific outcome as the cue [4,15,18–20]. For instance, a cue associated with chocolate selectively increases responding to actions that yield chocolate, but not to actions that yield a different snack. Specific PIT requires encoding the transition from the cue to the expected outcome to encode its identity and therefore is thought to reflect model-based processes. Conversely, general PIT, which reflects only the general motivational value of a cue, is considered to rely on model-free mechanisms [4,18].

Behavioral evidence supports the distinction between specific and general PIT being model-based versus model-free. Both studies with rodents [18,21,22] and humans [23] indicate that specific PIT, but not general PIT, is sensitive to outcome devaluation, suggesting that specific transfer may be more model-based. However, in humans, these effects are less consistent, possibly because explicit knowledge about task contingencies can influence behavior during the transfer phase, as such knowledge may allow participants to strategically adjust their responses following outcome devaluation, thereby obscuring the computational processes underlying PIT [17,18,24]. Together, this supports the view that PIT recruits multiple computational systems, both model-free and model-based.

PIT effects are typically assessed through separate instrumental and Pavlovian training phases, followed by a transfer phase in which Pavlovian cues are presented during instrumental responding under nominal extinction. A diverse set of paradigms has been employed in the literature, using both appetitive and aversive conditioning. The full transfer paradigm captures both general and specific PIT [19], whereas the specific transfer design isolates only specific PIT [25], and the general transfer paradigm isolates general PIT alone [26]. All of these paradigms use multiple response options (i.e., multiple levers / response buttons). However, there is also a so-called single-lever paradigm [18,22,27]. This paradigm is simple, efficient, easily combined with other task structures, and widely used in studies of addiction and other mental illnesses, but it cannot distinguish between general and specific PIT [17,18]. Rat lesion studies suggest that single-lever PIT is impaired by lesions in areas associated with general but not specific PIT [18]. Moreover, in humans, single-lever PIT has been linked to model-free instrumental reinforcement learning measured by the two-step task [13]. Together, these findings are commonly interpreted as single-lever PIT being model-free. However, it remains unclear whether single-lever PIT is exclusively model-free, or whether it can also reflect model-based strategies. Addressing this issue is a central aim of the present study and, in addition to its widespread use, motivated our choice of this paradigm for our task.

As stated above, findings from PIT provide converging evidence for a dual-system view of Pavlovian responding. However, unlike instrumental learning, where paradigms like the two-step Markov decision task exist [7] are able to also dissociate trial-by-trial model-free and model-based learning systems computationally, no corresponding behavioral task exists for Pavlovian responding and more specifically for PIT. To address this critical gap, we developed a trial-by-trial two-stage Pavlovian learning paradigm designed to computationally dissociate model-free and model-based Pavlovian responding.

In research using the instrumental two-step task [7], it has been shown that intensive task instructions lead to increased reliance on model-based decision-making, with negligible contributions from model-free strategies [28]. This aligns with the notion that model-free and model-based systems compete for behavioral control depending on environmental uncertainty [1,29] and suggests that providing explicit instructions can reduce model uncertainty, thereby enhancing model-based engagement. Here, we sought to investigate whether single-lever PIT, which is commonly viewed as model-free learning, can also be influenced by model-based processes. Thus, we provided participants with detailed, repeated instructions along with comprehension checks before the task. This provided a direct test of whether single-lever PIT can reflect model-based processes, beyond influences of model-free learning under ideal circumstances of model certainty and motivation to act model-based.

*Mind wandering* is broadly defined as the shift of attention away from a primary task toward internally generated thoughts, i.e., it reflects a decoupling of attention from external input, favoring internal mentation [30,31]. Model-based learning, in contrast to the computationally simpler model-free learning, relies on constructing a detailed internal model of the environment and thus depends on cognitive functions such as working memory and processing speed [32,33]. When mind wandering occurs, these resources may be unavailable for model-based learning, thereby impairing its control of behavior, while model-free learning should remain unaffected. Support for this prediction was found using the instrumental two-step task [7] as mind wandering selectively impaired model-based but not model-free learning [34]. We therefore hypothesized that in our PIT task, mind wandering would disrupt model-based but not model-free learning, which would validate our operationalization of model-based and model-free PIT.

## Results

Our novel two-stage PIT task is illustrated in Fig. 1. First, during instrumental training (Fig. 1A), participants performed approach responses (via repeated button presses) or non-approach responses (by withholding responding) to different visual cues (playing cards). They learned via trial and error from probabilistic monetary feedback to approach and collect some cards and to non-approach and leave others. Afterwards, in the second part of the task, participants performed 480 trials of interleaved Pavlovian learning, PIT, and value query responses. Specifically, each individual trial of Pavlovian learning was immediately followed by a PIT trial and by a value query trial, where the order of the PIT and value query trials was counterbalanced across trials (i.e., either Pavlovian learning – PIT – query, or Pavlovian learning – query – PIT; Fig. 1B).

**Fig 1.**
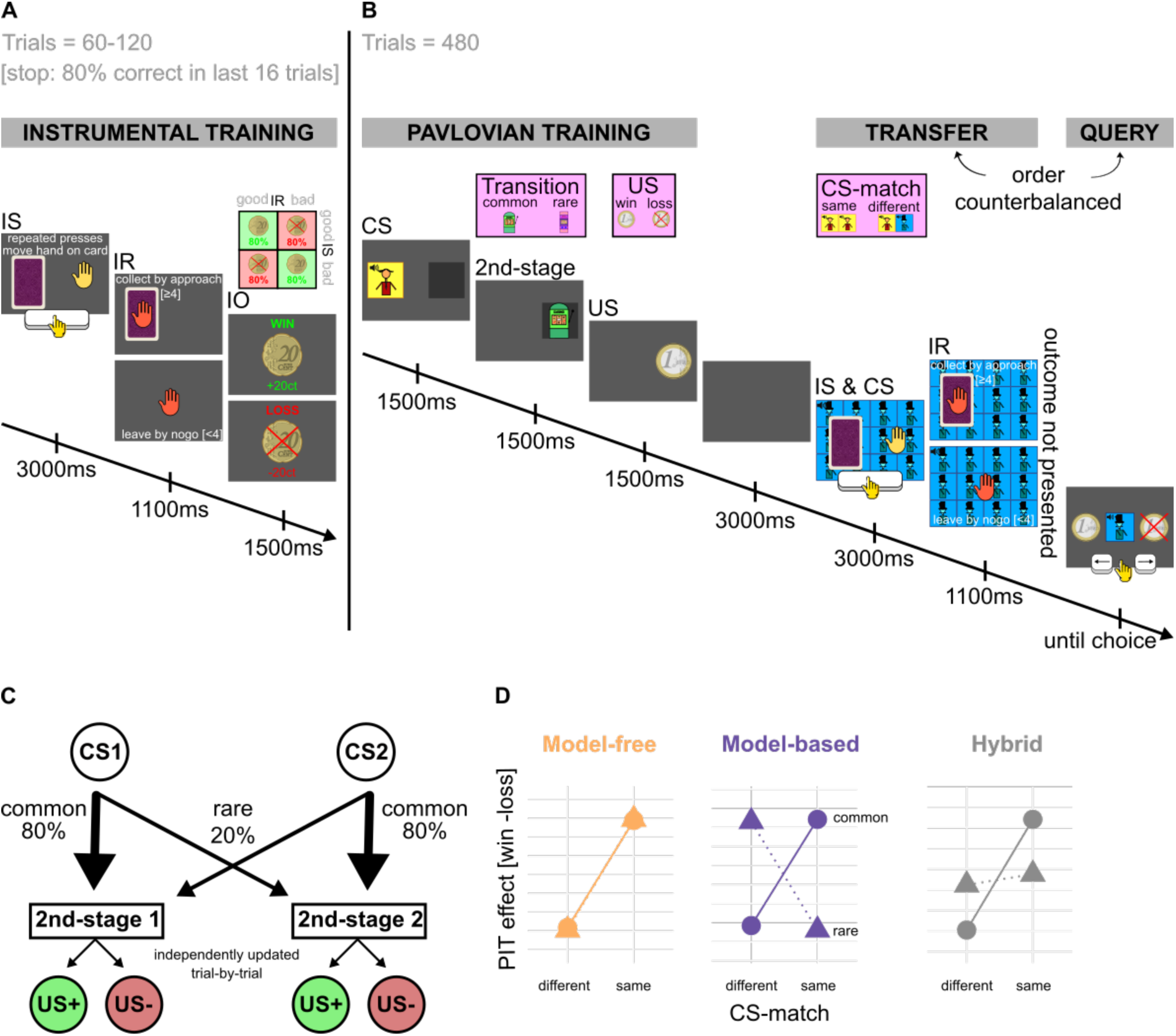
Two-stage Pavlovian-instrumental transfer (PIT) task. (A) Instrumental training. Participants first completed instrumental training: They learned from trial and error to collect “good” playing cards (instrumental stimuli, IS) by repeatedly pressing the spacebar (“go” instrumental response, IR) and to avoid “bad” cards by withholding responses, to maximize monetary wins and losses (instrumental outcome, IO). (B) Pavlovian training, transfer, and query. Participants completed Pavlovian training trials, interleaved with PIT transfer trials and forced-choice queries. The Pavlovian training exhibited a two-stage task structure. Each Pavlovian trial began with the appearance of one of two casino workers (conditioned stimuli, CS, 1^st^-stage), who led participants to one of two slot machines (2^nd^-stage) that yielded a monetary outcome of either winning or losing €1 (unconditioned stimulus, US). Each 1^st^-stage CS (casino worker) was associated with a "favorite" slot machine (2^nd^-stage), leading there 80% of the time (common transition condition), and to the alternative slot machine 20% of the time (rare transition condition). After a brief delay, participants completed a Pavlovian-instrumental transfer (PIT) trial, namely an instrumental trial under nominal extinction while a CS was shown in the background. Then they responded to a forced-choice question asking whether that CS was currently more associated with winning or losing. The order of transfer and query trials was counterbalanced across trials. CSs could either match (same CS; not shown) or not match (different CSs; see figure) between Pavlovian training and transfer/query phases (CS-match condition). Every 60 trials, participants rated their current mind-wandering on a continuous scale from –1 (“attention completely focused on the task”) to 1 (“attention completely focused on other thoughts”). (C) Pavlovian transition structure. Each CS (1^st^-stage) predominantly transitioned to a specific 2^nd^-stage with 80% probability, which then led to a probabilistic win or loss. Reward probabilities for the 2^nd^-stage states (slot machines) followed Gaussian random walks with reflecting boundaries, leading to gradually changing win/loss probabilities. (D) Simulated computational model predictions. Behavioral responses during the transfer phase were simulated using model-free, model-based, and hybrid reinforcement learning systems. Plotted is the difference in response rate between win versus loss trials (i.e., the PIT effect) as a function of CS-match (same vs. different) and transition type (common vs. rare). The model-free system predicts that the PIT effect depends only on CS-match, reflecting direct reinforcement of the presented CS (but not the other/different, i.e., non-presented CS) regardless of transition type. The model-based system predicts an interaction between CS-match and transition probability, with a reversal of the size of the PIT effect for rare compared with common transitions. The hybrid model combines both effects, capturing contributions from model-free and model-based learning.

The Pavlovian training phase contained two task stages. The 1^st^-stage involved presentation of one of two audio-visual 1^st^-stage conditioned stimuli (CS; one of two different waiters in a casino, saying “hello” to the participant). 1^st^-stage CSs predicted probabilistic transitions to one of two 2^nd^-stage states (i.e., one of two slot machines), which in turn lead to probabilistic wins and losses (Fig. 1B+C). Win probabilities of slots machines slowly changed across trials according to bounded Gaussian random walks. Each 1^st^-stage CS predicted one 2^nd^-stage state with higher probability (80%, common transition), and the other with lower probability (20%, rare transition; Fig. 1C).

For the following PIT and value query, the 1^st^-stage CS was either the same shown in the corresponding Pavlovian conditioning trial, or a different one (condition CS-match). In each PIT trial, participants performed the instrumental task (i.e. selecting or not selecting a card), while a 1^st^-stage CS was presented in the background. No outcomes were shown, but participants were instructed that outcomes would count towards their balance (nominal extinction). Approach responses during PIT served as an implicit measure of Pavlovian value learning, with the expectation that win-predictive CSs would increase approach responses and loss-predictive CSs would decrease approach responses, reflecting a PIT effect. In value query trials, participants were instructed to select whether the given 1^st^-stage CS was currently more associated with a win or a loss. Correct responses in PIT (collecting good cards or leaving bad cards) and the value query (correctly telling whether or not the 1^st^-stage CS is currently more associated with win or loss) increased a performance-based bonus, motivating engagement and enabling trial-wise assessment of belief updating. Participants did not obtain trial-by-trial feedback on their bonuses to avoid further learning.

Using this trial-by-trial, two-stage Pavlovian learning design, we aimed to distinguish model-free, model-based, and hybrid learning strategies by leveraging their distinct computational signatures. In model-free learning, stimulus-outcome associations should be updated based solely on direct reinforcement history, i.e., exclusively for the actually presented 1^st^-stage CS but not for the other, non-presented 1^st^-stage CS, and thus ignoring the probabilistic transitions within the task. In contrast, model-based learning incorporates the transition structure between stages, enabling flexible inference about the relationship between cues and outcomes, i.e., inferring value updates also for the 1^st^-stage CS that was not presented in the current trial. In hybrid learning, PIT responding reflects a weighted mixture of model-free and model-based value updates, with the balance determined by the relative weighting of model-based and model-free contributions.

To derive model predictions for the PIT effects, we simulated model-free and model-based computational RL models (Fig. 1D). The model-free system predicted a PIT effect (i.e., increase in the number of button presses after a win compared to a loss) when the CS shown during PIT was the same as during Pavlovian training, but not when the different CS was presented, reflecting a main effect of CS-match on the PIT effect. By contrast, the model-based system predicted an interaction of transition frequency times and CS-match (Fig. 1D). The hybrid model predicted the presence of both a main effect and an interaction. Based on these results, we used the main effect of CS-match on the PIT effect as a behavioral marker of model-free PIT and the interaction CS-match and transition type as a behavioral marker of model-based PIT. Moreover, we applied the same logic of model-free and model-based behavioral markers to the analysis of the forced-choice query data.

We determined sample size using Bayesian sequential testing [35–37]. We computed null hypothesis Bayes factor tests for behavioral markers of model-free versus model-based learning on PIT responses, with boundaries of 1/6 < BF_10_ < 6 (for details see methods section). Bayesian sequential testing led to N = 71 participants (details see below).

### Behavioral Analysis: Forced-choice Query

We first assessed whether participants understood the task structure, specifically, how to infer current win/loss probabilities for the CS, by analyzing responses in the trial-by-trial forced-choice queries. On average, participants chose the response implied by the preceding Pavlovian conditioning trial in 58% of trials (*SD* = 14%; across all participants, 19778 of 34080 trials were successes), exceeding chance level (*p₀* = 0.5, exact binomial test, *p* < 0.001) and aligning with performance reported in comparable two-stage paradigms [7,34,38].

Next, we quantified model-based and model-free effects on query responses. As shown in Fig. 2A, responses followed a model-based pattern: the difference in query responses between win versus loss trials was influenced by an interaction of CS-match and transition type, yielding a robust model-based effect (*b* = 2.56, *SE* = 0.32; *BF_10_* = 4.89 × 10^10^; *Wilcoxon W_1-tailed_* = 2496, *p* < 0.001). In contrast, there was evidence for no main effect of CS-match, reflecting absence of model-free updating (*b* = –0.45, *SE* = 0.16; *BF_10_* = 0.01; *Wilcoxon W_1-tailed_* = 1174; *p* = 0.725). These findings indicate that participants relied on structured, transition-aware inference rather than direct reinforcement of the presented CS when making their query decisions.

**Fig 2.**
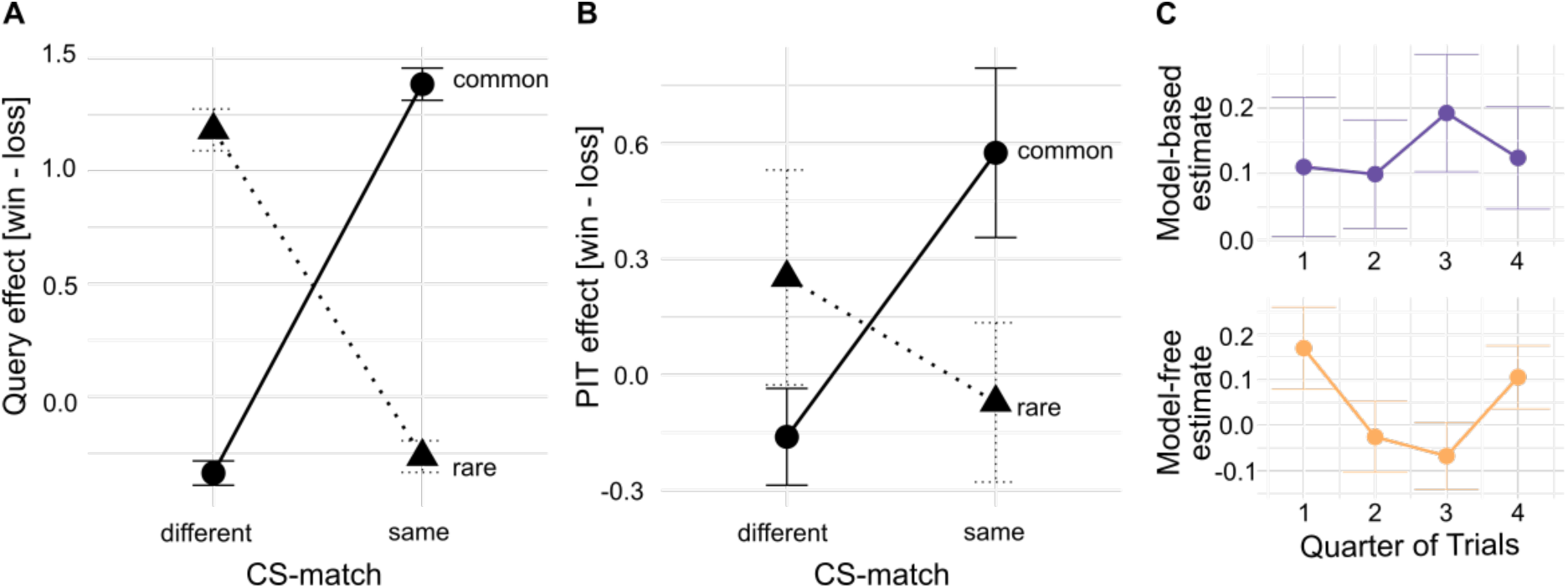
Query and two-stage Pavlovian-instrumental transfer (PIT) (A) Query behavioral results. During the forced-choice query trials, participants indicated whether a presented conditioned stimulus (CS) was currently more strongly associated with a win [coded as 1] or a loss [coded as –1]. Depicted is the average response difference between trials with a win versus a loss experienced in the preceding Pavlovian training trial. Responses are shown separately for CS-match (same vs. different) and transition type during Pavlovian training (common vs. rare). Error bars indicate the standard errors of the mean (SEM). The results show clear evidence for a CS-match x transition type interactional effects on the query effect, reflecting model-based responding. (B) PIT behavioral results. The PIT effect reflects the difference in response rate (number of button presses) for trials with a win versus a loss in the preceding Pavlovian training. The PIT effect is shown separately for CS-match (same vs. different) and transition type (common vs. rare). Error bars are SEM. PIT effects also exhibit an interaction CS-match x transition type, reflecting model-based responding. (C) Model-based and model-free estimates across task stages, shown for each quarter of trials.

### Behavioral Analysis: PIT

Critically, we next examined responding during the transfer phase. Consistent with the query results, the PIT effect also followed a model-based pattern, though less strongly (Fig. 2B). Bayes factors provided evidence that the PIT effect was modulated by an interaction of CS-match and transition frequency (Bayesian sequential sampling based on individual effect estimates: *b* = 0.13; *SE* = 0.06; *BF_10_* = 6.42; *Wilcoxon W_1-tailed_* = 1689, *p* = 0.009; hierarchical model estimate for full sample: *BF_10_* = 26.15). This supports model-based control of PIT responding, where participants adjust their responses based on inferred state-outcome relationships rather than simple reinforcement history. At the same time, Bayes factors were inconclusive with respect to whether the main effect of CS-match modulated the PIT effect (Bayesian sequential sampling based on individual effect estimates: *b* = 0.04, *SE* = 0.04; *BF_10_* = 1.02; *Wilcoxon W_1-tailed_* = 1445, *p* = 0.17; hierarchical model estimate for full sample: *BF_10_* = 4.44), providing no clear evidence for or against model-free PIT. These findings suggest that response behavior during PIT was primarily modulated by Pavlovian values estimated from structured, transition-aware inference. To examine how these patterns evolved over the course of the task, Fig. 2C provides a closer descriptive view by visualizing model-based and model-free estimates across quarters of trials.

### Computational Modelling

To quantitatively assess participants’ learning strategies during PIT across trials, we fitted three computational models, model-free, model-based, and hybrid, to each participant’s data using expectation maximization [12,39]. All models assumed a Poisson distribution for PIT response rates and included a general intercept (β_0_) and a slope for instrumental stimulus type (β_IS_). They differed in how Pavlovian influences were represented: the model-free model implemented temporal difference learning [3] with learning rates at both stages (α_1_, α_2_), an eligibility trace parameter (λ), and a model-free weight (β_MF_). The model-based model used a transition-aware update consistent with the Bellman equation [40] with a 2^nd^-stage learning rate (α_2_) and a model-based weight (β_MB_). The hybrid model combined both strategies, including all learning-related parameters (α_1_, α_2_, λ) and both weighting parameters (β_MF_, β_MB_).

We compared model fits at the individual level using Bayesian Information Criterion (BIC) scores. The model-based computational model provided the best fit overall (Fig. 3A), yielding significantly lower BIC scores than both the hybrid model (*Wilcoxon W* = 1994, *p* < 0.001) and the model-free model (*Wilcoxon W* = 2281, *p* < 0.001). Moreover, the model-based model exhibited the best model-fit for the majority of participants (Fig. 3B; being the winning model for 52 of the 71 participants), reinforcing the interpretation that participants relied on inferred state-outcome mappings.

**Fig 3.**
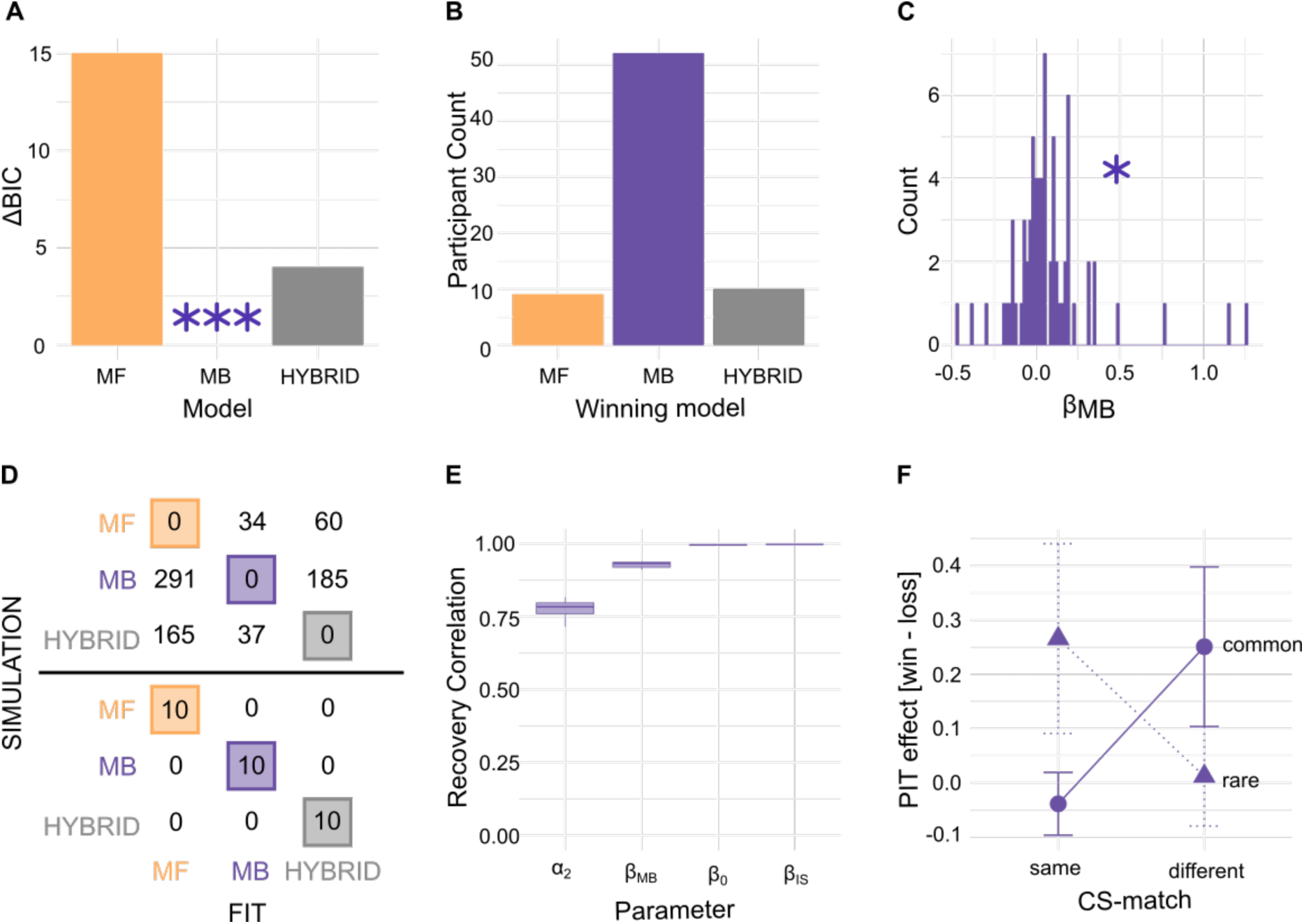
Computational model analysis. (A) Model comparison based on relative BIC scores (relative to best-fitting model). Across participants, the model-based model outperformed both the hybrid model (*W* = 1994, *p* < 0.001) and the model-free model (*W* = 2281, *p* < 0.001), suggesting evidence for the model-based strategy at the group level. (B) Winning model based on the BIC criterion for each participant. The model-based model best accounted for the majority of participants (52 out of 71). (C) Distribution of the model-based weighting parameter (β_MB_) estimated using the EM algorithm. The parameter estimate β_MB_ was significantly greater than zero (*p* < 0.05). Significance was assessed using MAP estimates with EM-derived priors but with the β_MB_ prior centered at zero. (D) Model recovery. Ten synthetic datasets were generated from each model and then refitted with all candidate models. The upper table displays the accumulated relative BIC values (relative to the best-fitting model) across refits, and the lower table shows how often each model was identified as the winning model. This analysis suggests that each model can be reliably distinguished based on its characteristic fit pattern. (E) Parameter recovery. For the winning model-based model, parameters were simulated from the fitted participants’ model estimates and re-estimated from the simulated data ten times. The correlations of recovered and true parameter values are depicted in the box plots. (F) Posterior predictive checks. The plot shows the behavioral pattern of the simulated data. The model reproduces key features of the empirical data, including the interaction of CS-match (same/different) and transition type (common/rare), supporting the adequacy of the model fit. (A + C) *, *p* < .05; **, p < .01; ***, p < .001.

For the winning model-based RL model, Fig. 3C depicts the distribution of parameter estimates for the model-based weight (β_MB_) across participants. To quantify the contribution of model-based control during PIT, we statistically confirmed that β_MB_ was significantly greater than zero (Wilcoxon *W* = 1715, *p* = 0.006), indicating a reliable reliance on transition-aware strategies. Model recovery, parameter recovery, and posterior predictive checks showed that the model was reliably identifiable and captured the central behavioral effects (Fig. 3D-F).

### Model-based Learning and Mind Wandering

To examine whether model-based control was modulated by participants’ attentional state, we assessed both state and trait mind wandering. To assess state mind wandering, after every 60 trials, participants rated their attentional focus during the preceding block on a continuous scale ranging from –1 (completely on-task) to 1 (completely off-task). We found that higher self-reported mind-wandering scores significantly predicted lower model-based behavioral estimates during those trial blocks (Fig. 4A; *b* = –0.34, *SE* = 0.17, *t*(160.76) = – 2.01, *p* = .024). Moreover, we assessed spontaneous and deliberate mind wandering using a questionnaire [34,41]. Individual differences in model-based PIT (behavioral marker) correlated negatively with deliberate mind-wandering trait scores (Spearman’s *ρ* = −0.27, *p* = .011) and, as a trend, also with spontaneous mind-wandering trait scores (Spearman’s *ρ* = −0.18, *p* = .071), suggesting that a greater dispositional tendency to disengage from task-relevant processing was associated with reduced model-based learning (Fig. 4B). In contrast, no measure of mind wandering was related to model-free signatures (*p* > 0.1).

**Fig 4.**
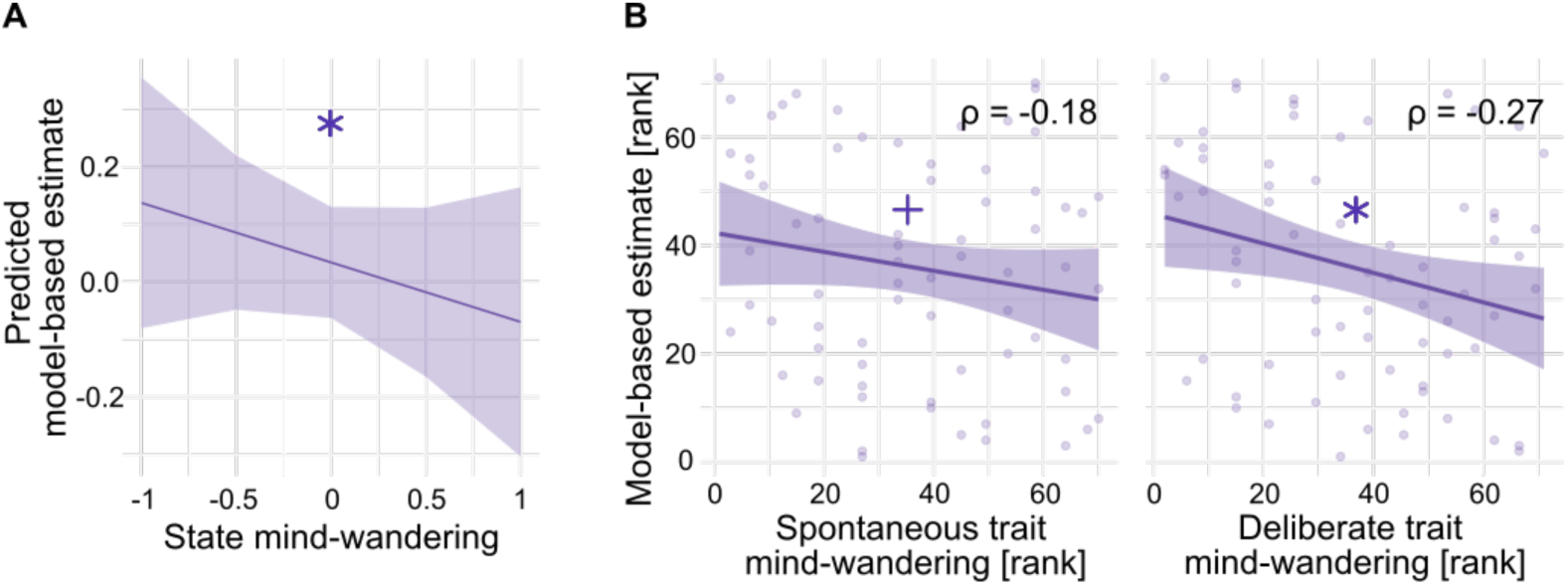
Relationship between model-based behavior and mind-wandering. (A) Association between self-reported state mind wandering and model-based estimates. Every 60 trials, participants rated their attentional focus during the preceding block on a continuous scale from –1 (“attention completely focused on the task”) to 1 (“attention completely focused on other thoughts”). Higher mind-wandering ratings predicted reduced model-based estimates across blocks (*b* = –0.34, *SE* = 0.17, *t*(160.76) = –2.01, *p* = .024). (B) Associations between trait mind wandering and model-based estimates. Participants’ overall model-based estimates were negatively correlated with deliberate mind-wandering scores (Spearman’s *ρ* = –.27, *p* = .011), and a negative trend was observed for spontaneous mind-wandering (Spearman’s *ρ* = –.18, *p* = .071). Lines represent linear fits with 95 % confidence intervals; points denote individual participants (ranks). (A + B) ^+^, *p* < .10; *, *p* < .05.

## Discussion

In this study, we introduce a trial-by-trial two-stage Pavlovian instrumental transfer (PIT) paradigm designed to distinguish model-free versus model-based Pavlovian responding, using both behavioral and computational measures. Using this paradigm, we provided evidence that Pavlovian influences on instrumental behavior are not merely the product of model-free values but may instead be governed by model-based inference over structured task contingencies.

At the behavioral level, to probe explicit learning, we performed direct value queries in each trial, which revealed that participants relied on behavioral markers of model-based learning, demonstrating both an understanding of the task structure and a capacity to update beliefs about stimulus values. We also studied how trial-by-trial Pavlovian CS value elicited biases in ongoing approach behavior during PIT. We found that Pavlovian values were dynamically updated over time. Crucially, behavioral PIT responses were modulated by behavioral makers of model-based learning, as they were shaped not only by the value of the last US (win versus loss) and whether the CS was presented in the last Pavlovian conditioning trial (i.e., CS-match), but also by the transition structure linking cues to outcomes (common versus rare transitions). This pattern aligns with the notion of model-based control, where decisions are guided by an internal model of the environment’s structure, including state transitions and outcome expectancies [1,2,5], supporting the engagement of model-based control in single-lever PIT and thus Pavlovian responding.

Computational modeling confirmed and extended these findings. We fitted computational RL models assuming either pure model-free, pure model-based, or hybrid control to the participants’ data. We found that the model-based control model fitted the data best, both when averaged across participants as well as individually in the majority of participants. In addition, the model-based weighting parameter (β_MB_) was substantially positive, demonstrating systematic model-based influences.

We assessed mind wandering to capture variance associated with fluctuating attention and to validate our measures of model-based learning. The results showed that higher reports of mind wandering during the task (thought probes) predicted lower model-based scores. Moreover, higher trait levels of both deliberate and spontaneous mind wandering were negatively correlated with model-based control. Model-free control was not associated with mind wandering reports. These findings are consistent with prior results [34], showing that mind-wandering specifically impairs model-based behavior in the instrumental two-step decision task but leaves model-free learning intact. This validates our interpretation that our markers of model-based PIT are indeed associated with cognitive resources theoretically believed to underlie model-based computations: Model-based strategies are known to depend on cognitive resources like working memory and executive function, which are reduced when attention drifts away from the task, leading to impaired model-based control [32,33]. Our findings extend these results from instrumental learning to the Pavlovian domain, demonstrating that (model-based) PIT relies on fluctuating internal attentional state.

Based on the idea that specific and general PIT reflect model-based versus model-free PIT responding, clear hypotheses exist that model-free and model-based PIT effects may partly rely on distinct neural structures. In rodents, lesion and inactivation studies [19,42], as well as disconnection studies [43], indicate that specific transfer involves the basolateral amygdala and nucleus accumbens shell, whereas general transfer engages the central amygdala and nucleus accumbens core. In humans, fMRI studies have correspondingly implicated the amygdala and ventral striatum in PIT [44–46], and high-resolution fMRI could even support a neuronal dissociation in humans, with specific transfer associated with the ventral amygdala in the basolateral complex and the ventrolateral putamen, and general transfer linked to the dorsal amygdala in the centromedial complex [15]. Additionally, it was demonstrated that, in humans, specific, but not general, transfer depends on lateral prefrontal cortex activity in a transcranial direct current stimulation study [47]. Thus, it would be interesting to investigate these neural substrates during the two-stage Pavlovian-instrumental transfer task to assess whether model-free versus model-based behavior in this task engages and depends on similar areas.

The current study contributes to the growing field that emphasizes more individualized and nuanced perspectives on fundamental learning processes such as Pavlovian learning, enabling deeper investigation into their underlying mechanisms. Such insights are also crucial for advancing our understanding of psychological disorders and for improving and tailoring treatment approaches. Dysregulated and maladaptive Pavlovian learning is indicated in a variety of mental disorders like substance use disorders (SUDs) [14,48–52], impulsive disorders such as gambling disorder, compulsive sexual behavior, kleptomania, and gaming addiction [53,54], eating disorders, specifically emotional eating [55], obsessive-compulsive disorders (OCD) [56], anxiety disorders [57], mood disorders such as depression [58], and even schizophrenia [59,60]. This underscores the importance of investigating the mechanisms underlying learning processes in greater depth, as doing so may enable a more systematic targeting of individual needs in therapy. Moreover, it remains to be seen in how far the use of model-free versus model-based Pavlovian responding represents fixed traits or can be modified to alleviate clinical symptoms [61].

There are some limitations of our work. While we interpret our results primarily within a model-based versus model-free framework, it is important to note that this dichotomy may be overly simplistic. Recent work suggests that behavior traditionally labeled as "model-based" may also be explained by alternative, simpler strategies, such as the successor representation (SR) [62], which blends elements of model-free and model-based control by caching multi-step predictions of future states [63]. This may also explain our current findings on model-based PIT. Conversely, "model-free" behavior may not always reflect a purely value-cached strategy. Instead, participants might rely on "moldy models", partial, noisy, or misinformed internal models that still incorporate structural elements of the environment but fail to fully support flexible planning [64]. In our study, we performed additional debriefing analyses (details not reported) showing that some participants lacked a full, explicit understanding of the task structure even after completing it, which may have contributed to apparent deviations from purely model-based or model-free patterns for these participants. Future work should develop experimental paradigms allowing model comparison frameworks that test a wider range of possible strategies, including SR and partially informed models, and combine behavioral data with neural and more elaborate and varied debriefing measures to better characterize individual differences in strategy use.

We did not find clear evidence for model-free control in our PIT task. Instead, the Bayes factor evidence for behavioral markers of model-free PIT was undecided. This may be due to the limited sample size, which did not enable clear conclusions with respect to model-free behavior. However, the computational model comparisons supported a pure model-based model without a model-free component. While model-based markers were clearly present, it may still be possible that model-free strategies show up in our task design. Importantly, we had designed our task to maximize model-based and minimize model-free contributions. For that, we included detailed instructions and comprehension checks to increase correct model knowledge and reduce uncertainty, as well as a forced-choice query explicitly aimed at motivating model-based behavior [1,28,29]. Therefore, it seems possible that model-free PIT control is more prominent when task instructions are less detailed, leading to potentially weaker model-based and stronger model-free effects. This interpretation aligns with findings suggesting that model-free strategies can persist despite improved instruction, as seen in the two-step Markov decision task, where they exert a robust and somewhat automatic influence on behavior [65]. Future research should re-examine the potential contribution of model-free effects using larger sample sizes and designs specifically powered to detect model-free contributions.

In summary, this study contributes to a body of evidence indicating that Pavlovian responding is not solely governed by model-free mechanisms. The use of a trial-by-trial, two-stage PIT paradigm allowed for fine-grained assessment of different learning strategies and opens new directions for understanding the conditions under which model-based Pavlovian systems operate. With that, we were able to show that during a single-lever PIT task, participants can actively engage in Pavlovian inferences that are more complex than mere model-free updating for experienced CSs, and that this process is sensitive to both environmental structure and internal cognitive state. These findings refine our understanding of the computational architecture of Pavlovian systems and suggest that even reflexive-seeming behaviors can be shaped by deliberative processes.

## Methods

The study hypotheses, computational workflows, and analysis procedures, including the Bayesian sequential testing plan and stopping rules, were preregistered on OSF prior to data collection (https://osf.io/cyekz). Note that we slightly deviated from the preregistered sampling plan. We had preregistered to collect data until we found evidence for or against (a) model-based PIT and (b) model-free PIT. However, given the number of 71 participants, we decided to stop sequential testing once the data provided Bayes factor evidence for model-based PIT. Contrary to frequentist analysis, such changes in plan are compatible with Bayesian (sequential) hypothesis testing [35–37].

## Participants

Seventy-one healthy university students (49 females, 21 males, 1 diverse; age range: 19–35 years, *M* = 23.3, *SD* = 3.5) participated in the study. All participants provided written informed consent before participation. The task took about three hours (including breaks) and participants received between three and seven hours of partial course credit as compensation, from which four hours were contingent on task performance. Performance was calculated based on monetary wins and losses during the instrumental and Pavlovian training phases as well as the transfer phase, in addition to accuracy on value queries derived from the instructed model. Participants performing at or below chance received no bonus, those with average or above-average performance received the full four bonus hours, and participants falling in between were granted a partial bonus. The study was approved by the ethics committee of the HMU Health and Medical University Potsdam, and all procedures adhered to the Declaration of Helsinki.

### Prior Analysis and Sequential Bayesian Testing

To inform sample size and analysis, we simulated behavioral data a priori from a dual-control computational model combining model-free temporal difference (TD) learning (Sutton et al., 1998) and model-based learning consistent with the Bellman equation (Bellman, 1957). The model included parameters for the learning rates for the 1^st^- (α_1_) and 2^nd^-stage (α_2_), for eligibility traces (λ), for model-based vs. model-free weighting (ω), an intercept for instrumental response (β₀), and for a slope capturing the instrumental stimulus type effect (β_IS_). Using 2-step parameter estimates drawn from distributions informed by Schad et al. [33] and PIT parameter by Schad et al. [16], we simulated 100,000 agents performing 480 trials each under three learning scenarios: purely model-free (ω = 0), purely model-based (ω = 1), and hybrid (ω from empirical estimates).

Simulations confirmed that model-free control produces a main effect of CS-match on the PIT effect, while model-based control yields an interaction of CS-match × transition type (Fig. 1C). These effects were used to define statistical markers of learning mechanisms in empirical data.

To determine sample size, we implemented Bayesian sequential testing [35–37]. For each participant, we fit a linear model predicting PIT responses (i.e., number of button presses during transfer) using IS (go = 1 vs. nogo = −1), CS-match (same = 1 vs. different = - 1 in Pavlovian training and transfer), US (win = 1 vs. loss = −1), and transition type (common = 1 vs. rare = −1), extracting coefficients for the two-way interaction of CS-match and US and three-way interaction CS-match, transition, and US. We operationalized model-free responding as a two-way interaction of CS-match x US and model-based responding as a three-way interaction of CS-match x US x transition type. Note that, above (Fig. 1), we had defined both in terms of their interaction with the PIT-effect, coded as the difference between win and loss trials, i.e., the effect of CS-value, for easier understanding and visualization. We tested whether across all participants these interaction estimates were different from zero using a Bayes factor t-test (Savage-Dickey method) with priors for the effect sizes und the alternative hypotheses derived from the computational simulations. Data collection was preregistered to continue at least until 30 participants were collected and until sufficient evidence (1/6 > BF10 > 6) for the key interaction effects reflecting model-based and model-free PIT was obtained. Here, given the sample size of n=71 participants, we decided to diverge from the preregistered analysis plan and stopped data collection after evidence for model-based PIT was obtained. This was done because (a) we considered evidence in favor of model-based PIT to be crucial news for itself and (b) because we were unsure whether clear evidence for or against *both* effects simultaneously could be reached realistically with a reachable number of participants. Note that such changes in plan are valid in Bayesian (sequential) testing. Analysis was performed whenever 5 new participants were collected (Fig. 5). Sampling was stopped after 71 participants.

**Fig 5.**
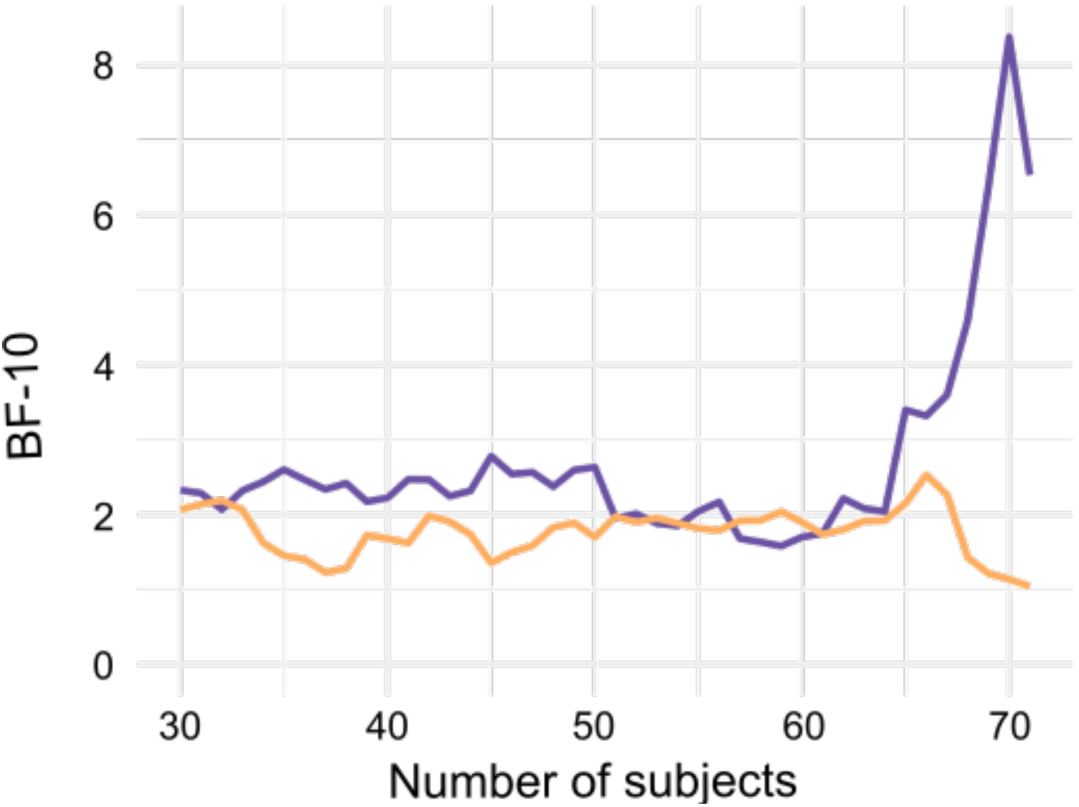
Bayesian Sequential Sampling. The sample size was not predetermined. Instead, we employed a Bayesian sequential sampling approach to determine when sufficient evidence had been accumulated. Specifically, we computed individual estimates of model-free behavior (defined as the main effect of CS-match on the PIT effect) and model-based behavior (defined as the two-way interaction between CS-match and transition type on the PIT effect) for each participant. At each interim checkpoint of five new participants, we calculated Bayes factors to assess the evidence for or against the null hypothesis that the estimates were greater than zero. Sequential monitoring began after data were collected from a minimum of 30 participants and continued until the Bayes factor for the model-based estimate exceeded 6, indicating strong evidence for a model-based contribution, or was smaller than 1/6, indicating evidence for the null hypothesis of no model-based contribution.

### Procedure

#### Task Overview

Participants completed a novel PIT paradigm (Fig. 1) designed to dissociate model-based and model-free Pavlovian learning influences on instrumental behavior. The task was programmed in MATLAB R2024a [66] using Psychophysics Toolbox Version 3 [67–69]. Participants were thoroughly instructed about the task and its underlying task structure (including transition probabilities between 1^st^- and 2^nd^-stage states) and were required to demonstrate sufficient understanding before beginning the experiment. Moreover, to promote active engagement and belief updating, participants responded to a direct query regarding stimulus-outcome associations, with accuracy linked to increased bonus payments.

#### Instrumental Training

Participants initially learned an instrumental card-collection game involving six distinct playing cards, presented randomly on the left or right side of the screen. Three cards required a “go” response (collecting the card) to maximize monetary gains, and three required a “no-go” response (leaving the card). Correct responses were reinforced with a 20-cent reward 80% of the time, introducing probabilistic reinforcement to encourage learning.

Each trial began with a card stimulus displayed for 3000 ms alongside a hand icon. Participants could “collect” the card by pressing the space bar multiple times (threshold: four presses, undisclosed to participants). If the threshold was met, the hand icon appeared over the card to indicate collection; otherwise, the hand appeared alone to indicate rejection. Response feedback was presented for 1100 ms. Outcome feedback, a 20-cent coin for reward or coin with a red cross for loss, followed for 1500 ms. Instrumental training lasted a minimum of 60 trials and continued until participants achieved a learning criterion of ≥80% correct responses over the last 16 trials or a maximum of 120 trials.

#### Pavlovian Training and Transfer Phase

Following instrumental training, participants completed a combined Pavlovian training and PIT phase structured as 480 repeated “casino visits.” Each trial involved three stages:

##### Pavlovian Training

Two abstract images of casino workers paired with a distinctive “hello” sound served as CS. Each CS probabilistically transitioned (80% common, 20% rare) to one of two slot machine images (2^nd^-stage), which independently produced monetary outcomes (win/loss of 1€) following a slowly drifting reward probability (random walk). The US was displayed after the 2^nd^-stage as a 1€ coin (win) or a 1€ coin with a red cross (loss). CS, 2^nd^-stage, and US were all presented for 1500ms.

##### Transfer

Participants performed the card-collection game from instrumental training, but with the background overlaid by multiple copies of the current CS accompanied by the “hello” sound, and under nominal extinction, i.e., with no outcome feedback.

##### Value Query

Participants indicated whether they believed the presented CS was more likely associated with a win or loss by choosing between corresponding icons on the screen using the arrows on the keyboard.

#### Mind-Wandering Assessment

Every 60 trials, participants rated their current attentional focus on a continuous visual analog scale ranging from “attention completely on-task” (–1) to “attention completely off-task” (+1). These ratings were used to assess the influence of state mind-wandering on model-based learning parameters [70]. Moreover, spontaneous and deliberate trait mind wandering was measured up front using a standardized questionnaire [41].

### Data Analysis

#### Behavioral Data

Statistical analyses were performed in RStudio v4.3.0 [71] and linear mixed-effects models were implemented using the lmerTest package [72]. To assess the extent to which behavior reflected model-based versus model-free learning, we analyzed participants’ responses as a function of three factors: US value (win vs. loss, capturing the PIT effect), transition type (common vs. rare), and CS-match (whether the CS presented during Pavlovian training matched the CS used during query and transfer).

The rationale was as follows: If a win (US) occurred during Pavlovian training and the CS led to its commonly associated 2^nd^-stage state, a model-based learner should infer that this CS predicts a valuable outcome. Accordingly, such learners would be more likely to respond “win” when queried and exhibit increased instrumental responding during transfer when the same CS is used. Following a rare transition, however, model-based reasoning would attribute the win to the alternate CS typically associated with the presented 2^nd^-stage state. Thus, the outcome should be discounted, and responses should reflect expectations based on prior knowledge rather than the most recent outcome. When CSs differed between Pavlovian training and the subsequent query and transfer, model-based reasoning predicts the opposite pattern, as outcome information should be mapped onto the CS most strongly linked to the updated 2^nd^-stage state.

In contrast, a model-free learner would assign value directly to the CS experienced during training, based solely on reinforcement history and independent of transition type. A win would reinforce the presented CS, leading to “win” responses during query and increased transfer responding when the same CS is presented. If a different CS is presented, the outcome should exert no influence, as model-free learning does not support inference about unexperienced stimuli or their associated states.

These predictions were tested during sequential sampling using regression models applied separately to each participant to obtain individual parameter estimates for subsequent group-level analyses. For the forced-choice query data, we fit logistic regression models with a logit link function to predict trial-by-trial responses (coded as win = 1, loss = – 1). Predictor variables included current US (win = 1, loss = –1, during the Pavlovian two-stage trial), CS-match (same = 1, different = 0), and transition type (common = 1, rare = –1), together with all interaction terms. For the PIT data, we fit linear regression models predicting trial-by-trial instrumental responding, using US (capturing PIT), CS-match, and transition type, again including all interactions, as well as including instrumental stimulus (IS)-type (go = 1, no-go = –1). For all models, predictors were coded using sum coding (–1/1) to maintain balanced contrasts [16]. Please note that for visualization, response measures were expressed as the difference between win and loss trials (i.e., using the PIT effect as the dependent variable), whereas statistical analyses included US-value (i.e., the PIT effect) as an explicit predictor term.

From each participant’s fitted statistical model for query and transfer responses, we extracted regression coefficients corresponding to the two-way interaction between CS-match and US-value, interpreted as the model-free index, and the three-way interaction among CS-match, US-value, and transition type, interpreted as the model-based index. These coefficients quantify, respectively, the degree to which behavior reflected direct reinforcement of the presented CS (model-free) and outcome updating modulated by transition structure (model-based).

Individual model-based and model-free coefficients were then entered into group-level analyses. We used a Bayesian approach to test whether each coefficient was greater than zero, applying the Savage–Dickey density ratio method. To obtain priors, we used the empirical distributions of the interaction effects obtained from 3,000 hybrid control model simulations. Specifically, the prior for the model-free term (US × CS-match) was set to a normal distribution with M = 0.05 and SD = 0.05, and the prior for the model-based term (US × CS-match × transition) was set to a normal distribution with M = 0.09 and SD = 0.09.

To confirm PIT results, we fitted a Bayesian hierarchical regression model using brms [73,74] to the full sample of all 71 participants. Transfer response rates were modeled with a Gaussian likelihood. Fixed effects included main effects of IS, US-value (capturing PIT), CS-match, and transition, as well as all interactions among US-value, CS-match, and transition. Subject-level random intercepts and random slopes were included for all predictors, with random-effect correlations omitted. Priors were specified using a simulation-informed procedure based on an a priori hybrid computational model. Synthetic datasets (3000 simulated subjects) were generated from a priori model predictions, and a frequentist mixed-effects model with an identical structure was fit to the simulated data. Fixed-effect estimates from this model were used to define weakly informative Normal priors centered at zero, with standard deviations equal to the absolute value of the corresponding estimates. Random-effect standard deviation priors were specified analogously. Priors with zero variance were replaced by Normal(0, 0.001) distributions to ensure numerical stability. Models were fit using four Markov chain Monte Carlo chains with 2,000 iterations per chain, including 500 warm-up iterations. Directional hypotheses indexing model-free and model-based learning were tested using posterior evidence ratios (Bayes factors), quantifying evidence that the US-value x CS-match (model-free) and CS-value x CS-match x transition (model-based) interactions were positive, with evidence ratios greater than 1 indicating increasing support.

#### Computational Model Fitting

We used computational modeling to distinguish model-free, model-based, and hybrid control processes in PIT. All computational modeling and simulation procedures were implemented in MATLAB [66]. To characterize the learning mechanisms underlying PIT, we fitted three reinforcement learning models, including model-free, model-based, and hybrid control, to each participant’s trial-by-trial PIT responses. Trial-by-trial response counts during the transfer phase were modeled as Poisson-distributed. Accordingly, Poisson log-likelihoods were computed for model fitting. Each model included a general intercept (β₀) and a slope parameter (β_IS_) to capture the influence of instrumental stimulus type (go vs. no-go) on response rate. The models differed only in their representation of the influence of effective Pavlovian value (V_pav_).

For model-free control, we implemented temporal difference learning (Sutton & Barto, 1998). This model included four additional free parameters (6 in total): the separate learning rates for the 1^st^- and 2^nd^-stage of the Pavlovian training (α_1_, α_2_), an eligibility trace decay parameter (λ), and a model-free weighting parameter (β_MF_) that modulated the influence of learned stimulus values on PIT behavior. For each trial (t), a prediction error, δ_1,t_, denoting the difference in value expectation between the presented 2^nd^-stage, V_2_(s_2,t_), and the presented 1^st^-stage, V_1_(s_1,t_), and a prediction error, δ_2,t_, denoting the difference between the outcome value (r_t_), i.e., the US in the present Pavlovian training, and the expected value based on the 2^nd^-stage, V_2_(s_2,t_), were calculated.

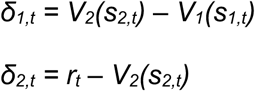

Based on both prediction errors and the learning rate for the 1^st^-stage, value expectations for the current 1^st^-stage, V_1_(s_1_,t), i.e., the CS in the present Pavlovian training, and the current 2^nd^-stage, V_2_(s_2_,t), were updated.

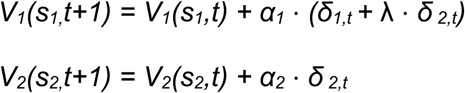

By incorporating the model-free weighting parameter into the newly informed 1^st^-stage value expectations, the combined effective Pavlovian value expectations are updated.

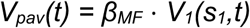

For the model-based control model, we employed a transition-aware, model-based updating mechanism aligned with the Bellman equation (Bellman, 1957). This model relied on two additional free parameters (4 in total): a 2^nd^-stage learning rate (α_2_) and a model-based weighting parameter (β_MB_). During every trial (t), a prediction error, δ_2,t_, representing the difference between the outcome value, r_t_, i.e., the US in the present Pavlovian training, and the expected value based on the presented 2^nd^-stage, V_2_(s_2_,t), was calculated.

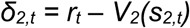

Based on the 2^nd^-stage’s prediction error (δ_2_) and the 2^nd^-stage learning rate (α_2_), the value expectation for the current 2^nd^-stage, V_2_(s_2_,t), was updated.

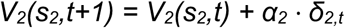

Afterwards, the instructed transition probabilities, which code the respective probabilities that each 1^st^-stage state leads to a given 2^nd^-stage state, were incorporated to generate a model-based value expectation update based on the current information for both possible 2^nd^-stages.

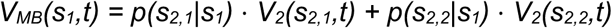

By incorporating the model-based weighting parameter, the combined effective Pavlovian value expectations were updated.

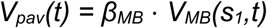

The hybrid model integrated both value systems, combining model-free and model-based value estimates accordingly, and contained all seven free parameters (β_0_, β_IS_, α_1_, α_2_, λ*, β_MB,_ β_MF_*).

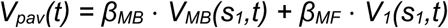

All models were fit using an expectation–maximization procedure (Dempster et al., 1977; Huys et al., 2011). In the estimation step (E-step), subject-level parameters were fitted by maximum a posteriori (MAP) [75,76] estimation, using MATLAB’s *fminunc* function for unconstrained nonlinear optimization. Parameters were transformed where necessary to respect natural bounds (learning rates and eligibility trace ∈ [0,1]; intercept, instrumental slope, and weighing parameters unbounded). Multiple random initializations and restarts were employed to avoid local minima. In the maximization step (M-step), group-level Gaussian priors for parameter means and variances were updated based on the current subject-level estimates. E- and M-steps were iterated until convergence, defined as a change of less than 0.01 in the MAP objective (log-likelihood plus prior means and standard deviations for all parameters). Across multiple EM runs, the solution with the highest penalized log-posterior was retained. Subject-level MAP estimates and Hessian-based uncertainty approximations were obtained from the EM procedure.

Model fits were evaluated using the Bayesian Information Criterion (BIC; Schwarz, 1978). For each participant, BIC was computed as

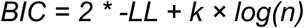

where *-LL* is the negative log-likelihood of the fitted model with optimized parameter estimates (MAP estimates based on the EM), *k* is the number of free parameters, and *n* is the number of observations for that participant. Lower BIC values indicate better model fit after accounting for model complexity and sample size. To compare models at the group level, we averaged individual BIC values across participants and tested differences between models using two-sided Wilcoxon signed-rank tests. Model identifiability was assessed via model recovery: we simulated datasets from each model using the estimated group-level parameters and re-fitted all candidate models to verify that BIC correctly identified the true data-generating model.

To assess the reliability of our parameter estimates and the adequacy of the winning model, we performed parameter recovery and posterior predictive checks. Specifically, we simulated 10 synthetic datasets per participant based on the model-based model and their individually estimated parameters. The simulated data were then re-fitted to recover parameters, which were compared to the original estimates (parameter recovery), and the simulated behavioral patterns were compared to the empirical data (posterior predictive checks).

Finally, we tested whether the weighting parameter for model-based control (β_MB_) was significantly greater than zero. Because the hierarchical EM procedure shrinks individual estimates toward the group mean, direct statistical testing on EM-derived parameters against zero may be biased. To address this, we performed an additional MAP estimation, using the EM-derived parameter distributions as priors but fixing the prior mean of the model-based weighting parameter (β_MB_) at zero. This approach ensured unbiased inference for β_MB_ while retaining prior information about parameter variability. The resulting participants’ estimates of β_MB_ were then tested against zero using a one-sided Wilcoxon signed-rank test.

#### Model-based Learning and Mind Wandering

Statistical analyses were performed in RStudio v4.3.0 [71] and linear mixed-effects models were implemented using the lme4 package [77]. To assess the relationship between state mind-wandering (thought probes) and model-based learning, linear mixed-effects models were fit with block-wise model-based estimates (interaction US x CS-match x transition type on response rate) as the dependent variable and predictors including momentary mind-wandering ratings and the time of the mind-wandering probe (i.e., block number). To evaluate the association between individual differences in mind-wandering traits and model-based learning, Spearman rank-order correlations were computed between questionnaire scores for spontaneous and deliberate mind-wandering and participants’ model-based learning indices. We conducted one-tailed tests with the a priori hypothesis that mind-wandering is negatively associated with PIT.

## Code and Data Availability

Anonymized data and all custom scripts used for data preprocessing, statistical analysis, computational modeling, and visualization are available at OSF (https://osf.io/mydvc/overview?view_only=52138a2f6c94428cba5eae31b7c54207).

## Acknowledgments

We are grateful to everyone who provided helpful input, discussions, and feedback. We are especially thankful to Charlotte Weber and Semra Merseli for their assistance with data collection. This work was supported by the Health and Medical University Potsdam and the Medical School Berlin through funding and the provision of laboratory space.

